# γBOriS: Identification of Origins of Replication in Gammaproteobacteria using Motif-based Machine Learning

**DOI:** 10.1101/597070

**Authors:** Theodor Sperlea, Lea Muth, Roman Martin, Christoph Weigel, Torsten Waldminghaus, Dominik Heider

**Affiliations:** Faculty of Mathematics and Computer Science, Philipps-Universität Marburg, D-35043 Marburg, Germany; Institute of Biotechnology, Faculty III, Technische Universität Berlin (TUB), Straße des 17. Juni 135, D-10623 Berlin, Germany; Chromosome Biology Group, LOEWE Center for Synthetic Microbiology (SYNMIKRO), Philipps-Universität Marburg, D-35043 Marburg, Germany

**Keywords:** origin of replication, machine learning, DNA classification, database, Gammaproteobacteria

## Abstract

The biology of bacterial cells is, in general, based on the information encoded on circular chromosomes. Regulation of chromosome replication is an essential process which mostly takes place at the origin of replication (*oriC*). Identification of high numbers of *oriC* is a prerequisite to enable systematic studies that could lead to insights of *oriC* functioning as well as novel drug targets for antibiotic development. Current methods for identyfing *oriC* sequences rely on chromosome-wide nucleotide disparities and are therefore limited to fully sequenced genomes, leaving a superabundance of genomic fragments unstudied. Here, we present *γ*BOriS (<u>Gamma</u>proteo<u>b</u>acterial *<u>ori</u>C* <u>S</u>earcher), which accurately identifies *oriC* sequences on gammaproteobacterial chromosomal fragments by employing motif-based DNA classification. Using *γ*BOriS, we created BOriS DB, which currently contains 25,827 *oriC* sequences from 1,217 species, thus making it the largest available database for *oriC* sequences to date.

## Introduction

Before every cell division, bacteria need to duplicate their genetic material in order to ensure that no information is lost. This essential process, called DNA replication, initiates in a highly regulated manner at specific chromosomal sites called *oriC* and is coordinated with many other cellular mechanisms (1, 2). Usually, bacteria contain (multiple compies of) a single chromosome and this chromosome contains a single *oriC* sequence, although there are exceptions as, e.g., Vibrionales contain two chromosomes (3, 4).

As many different proteins need to bind to and act upon *oriC* in order for initiation to occur, *oriC* contains many different protein binding sites and DNA motifs (5, 6). While there is a high level of variation between *oriC* sequences of different organisms, there are also some nearly universal features of *oriC* sequences (7–9). Among these are 9 bp short DNA motifs called DnaA boxes, which act as binding site for the initator protein DnaA, and AT-rich regions, where the DNA double helix unwinds before the replication machinery is loaded onto the DNA (10, 11). Furthermore, *oriC* contains binding sites for proteins that relay information on the status of the cell. Therefore, *oriC* sequences can be considered as biological information compiler and processors (12).

All currently available computational methods for the identification of *oriC* sequences in bacterial chromosomes rely on nucleotide disparities on the leading and lagging strand of the DNA double helix (13–17). As replication usually extends from *oriC* bidirectionally, it is one of two chromosomal sites where the leading and lagging strand switch places. The most frequently used disparity, the GC skew, usually assumes a V-or inverted V-shape with its minimum indicating the presence of the origin of replication (18, 19). However, due to natural variation, the shape of the skew can only reliably be asserted when analysing whole chromosomal sequences. Combining the GC skew with the location of DnaA boxes, Ori-Finder (20), was used to create the current state-of-the-art *oriC* database DoriC (21, 22).

While existing methods for the annotation of oriC sequences are mainly based on statistical approaches, motif-based approaches for DNA sequence classification by machine learning might be a promising alternative. Machine learning methods, in particular deep neural networks (CNNs) have been widely used already for similar tasks (23–29). However, these methods are notorious for needing big amounts of data and computing power. Support vector machines (SVMs) that perform classification on the basis of *k*-mer (i.e., *n*-gram) counts represent a less data-intensive alternative, and have even been shown to outperform CNNs when training data is small in number (30, 31). Some *k*-mer-SVMs even allow mismatches or gaps in these *k*-mers, leading to more realistic models of DNA motifs, which are subject to natural variation (32–34).

In the current study, we present *γ*BOriS (<u>Gamma</u>proteo-<u>b</u>acterial *<u>ori</u>C* <u>S</u>earcher), a tool that is able to identify *oriC* sequences for Gammaproteobacteria. This class of organisms contains many model organisms (e.g., *Escherichia coli, Vibrio cholerae* and *Pseudomonas putida*), and causative agents for serious illnesses such as such as cholera, plague and enteritis, which makes it a highly relevant study object. Making use of recent developments in the fields of DNA sequence classification and machine learning, *γ*BOriS enables *oriC* identification on both full chromosomes as well as chromosomal fragments, which drastically increases the number of sequences that can be searched for *oriC* sequences. Finally, using publicly available Gammaproteobacterial chromosomal fragments as input for *γ*BOriS, we gathered the largest dataset of bacterial *oriC* sequences available to date, BOriS DB.

## Materials and Methods

### A. Data curation and creation

A ground truth *oriC* dataset was compiled using a semi-automated method described in (9, 35, 36). A given chromosome is first split it into 2.5 kb fragments that are centered around intergenic regions and then, for those fragments close to the minimum of the chromosome’s cumulative GC-skew, their respective probability of unwinding is calculated using WebSIDD (37). Default values (37 °C, 0.1 M salt, circular DNA, copolymeric) were chosen for the predictions, and negative superhelicity values were tested in the range of *σ*.

A dataset of seed sequences was created by extracting the central 9 bp from *oriC* sequences in the ground truth dataset. Negative sequences for the initial classification training were collected by picking, for each chromosome present in the positive dataset, another sequence of the same size with the same seed sequence from the respective chromosome. In order to be able to identify the optimal fragment length for classification, the length of both the positive as well as the negative sequence were varied from 150 to 1500 bp in steps of 50 bp

For cutoff selection, a highly imbalanced dataset was created by extracting all fragments of a given length around each of the seed sequences from each of the chromosomes in the positive dataset.

Both the balanced as well as the imbalanced datasets were split into training and testing datasets using a 70%-30% split, leading to 318 chromosomes in the former and 141 chromosomes in the latter.

Chromosomes were downloaded from the NCBI refseq ftp server. For BOriS DB, a list of Refseq organisms was taken from ftp://ftp.ncbi.nlm.nih.gov/genomes/refseq/bacteria/assembly_summary.txt, a list of chromosomes for the UBA genomes was taken from Supplementary 2 of (38).

### B. Sequence classification using LS-GKM models

The support vector machines used as classifiers in this study derive distance matrices from a set of input sequences by counting substrings and comparing their numbers in sequence pairs directly, making these approaches faster and less memory intensive. The Spectrum Kernel is based on simple *k*-mer composition differences (30). However, LS-GKM and gkm-SVM models calculate differences between *k*-mers by allowing for mismatches and small differences between the *k*-mers (33, 39).

### C. *oriC* database comparisons

Comparisons were performed between BOriS DB v1 and DoriC v6.5, which were the latest accessible versions of the databases at the time of writing.

Pairs of sequences from different datasets were compared by calculating the length of the longest common substring and dividing it by the length of the shorter sequence. Two sequences were considered to be identical if the relative sequence identity was above a cutoff of 0.7. This cutoff was chosen in order to include overlapping sequences.

The internal consistency of the sequence datasets was evaluated by calculating all-vs-all sequence similarities from pairwise sequence alignments after making the sequences in the datasets of same length. Using multidimensional scaling and hierarchical clustering (as implemented in the Python packages scikit-learn and scipy, respectively (40, 41)), these distance matrices were visualized. A database was deemed more consistent if the degree of clustering is higher or if *oriC* sequences from closely related organisms are close on the tree.

## Results

### Implementation of *γ*BOriS

The stand-alone version of *γ*BOriS is implemented in R and requires a Linux operating system, whereas the frontend of the webserver is written in jQuery and can be used without any software requirements. As input file, *γ*BOriS takes a fasta-formatted file containing one or more DNA sequences of any length and returns two fasta-formatted text files: One contains fragments *γ*BOriS identified as *oriC* and the other contains DNA fragments for which the classifier abstained from a decision (see Methods). *γ*BOriS is composed of three modules that were adjusted for and trained on Gammaproteobacterial *oriC* sequences (fig. 1). The core module consists of a motif-based sequence SVM, whose parameters were chosen in order to maximize the AUC on discrimination of *oriC* from non-*oriC* sequences in a balanced dataset (see Methods, fig. 2). To this end, we trained a total of 12,877 LS-GKM and Spectrum Kernel SVMs (30, 32, 33) with varying parameters and sequence fragment sizes. The highest AUC of 0.977 on the test dataset, was achieved with a LS-GKM model trained with 1250 bp fragments, a word length of 10 bp with 6 informative columns and at most 4 mismatches (see supplementary information).

**Fig. 1.**
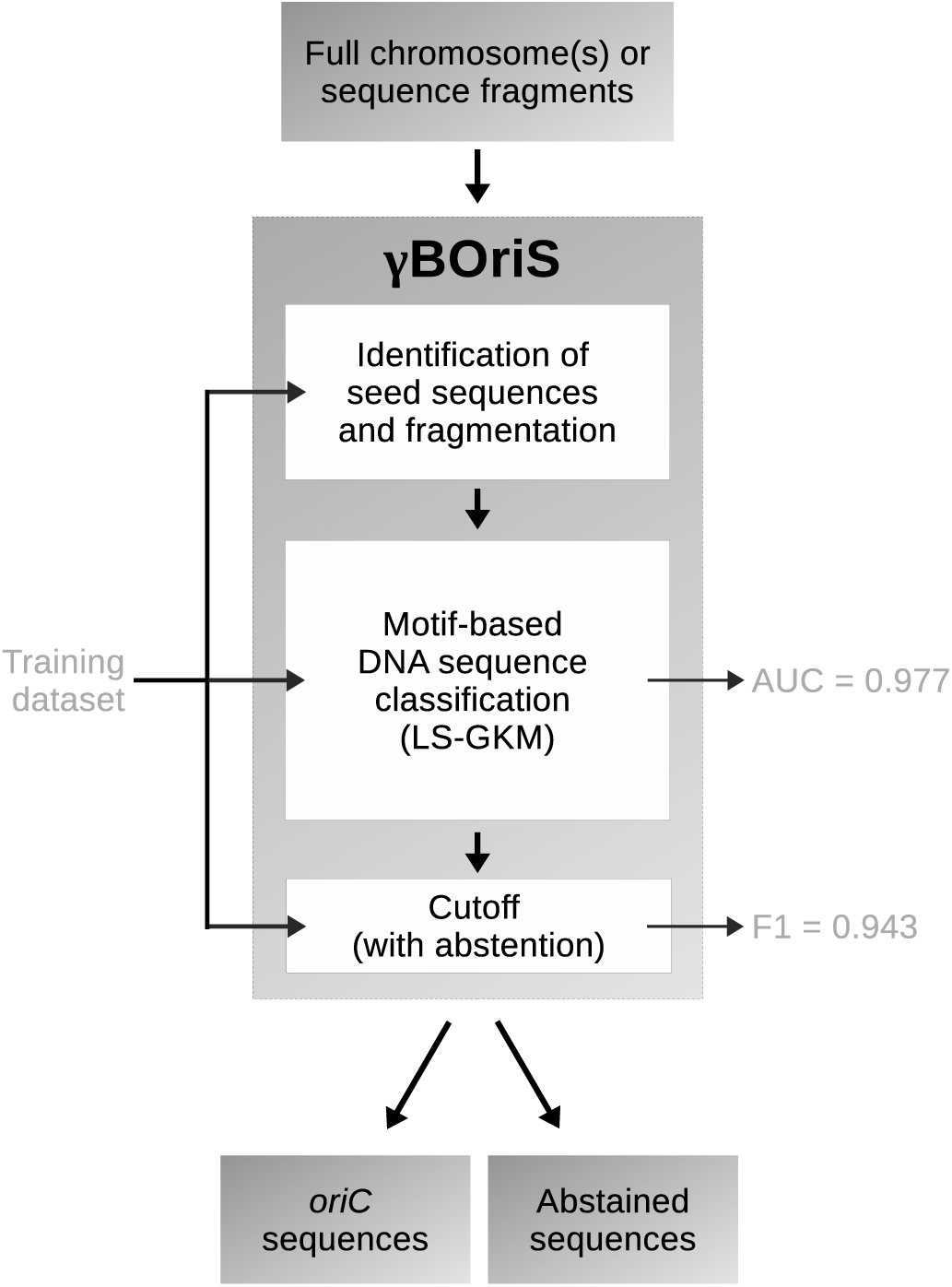
The structure of *γ*BOriS. Both the usage (top-down) as well as, schematically, the training process of the three modules and the training results as measured on a test dataset (left-to-right) is shown.

**Fig. 2.**
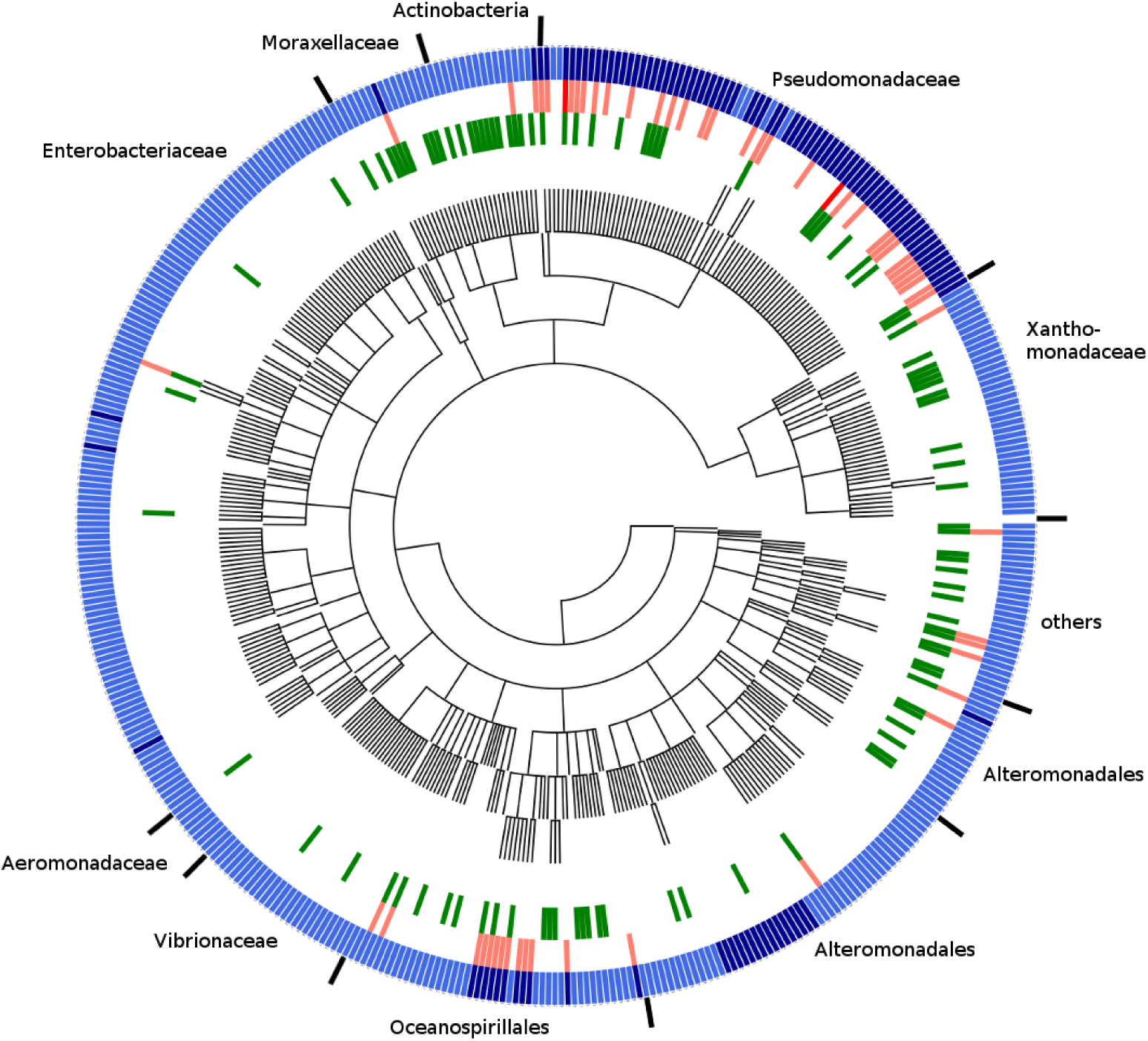
Taxonomic distribution of organisms, number of sequences and *γ*BOriS prediction results for the initial dataset. Outer ring: dark blue signifies chromosomes that contain two, light blue those that contain a single *oriC* sequences in the initial dataset. Middle Ring: Light red signifies chromosomes for which BOriS doesn’t recognize one of the *oriC* sequences, dark red signifies those for which two *oriC* sequences aren’t identified. Inner ring: Green represents chromosomes for which BOriS predicts falsely positive *oriC* sequences.

To turn this sequence classifier into a sequence identifier, the first module of *γ*BOriS splits the input sequence into a manageable number of candidate fragments by picking only fragments centered around an occurence of a so-called seed sequence. This list of seed sequences was created based on the initial *oriC* dataset by extracting the central 9bp sequences from the *oriC* sequences used for training of the classifier. This choice of seed sequences was validated by showing that all *oriC* sequences in the test dataset are centered around one of the seeds defined this way from sequences in the training dataset.

Finally, the third module of *γ*BOriS assigns a class label to every fragment based on the classification value obtained for this sequence in the second module. As, for one input sequence, the number of candidate sequences is expected to be much higher than the number of correct *oriC* sequences, this is a highly imbalanced problem. To mitigate high numbers of false positive classifications, we make use of the concept of classification with abstaining (42). To this end, two cutoffs are employed; below the lower cutoff, fragments are labeled “negative”, above the upper cutoff, fragments are labeled “positive” and between the cutoffs, the classifier abstaines from labeling the fragments. In the choice of cutoffs, we aim to both maximize the value of F1 and minimize the number of correct *oriC* sequences for which this module abstained from classification, leading to a Pareto-optimal state. For the sequences used to train *γ*BOriS, we found that normalizing the classification values of the fragments extracted from one sequence to a range between [0, 1] and employing cutoffs of 0.99 and 0.41 lead to the best result on the test dataset (F1 of 0.943 with 0.7% of correct *oriC*s in the abstained space). *γ*BOriS as a web tool as well as a stand-alone software and its source code are freely available at BOriS.heiderlab.de.

### Construction of BOriS DB

In order to create a large dataset for gammaproteobacterial *oriC* sequences, we applied *γ*BOriS to all chromosomes and chromosomal fragments present in the Refseq database (restricted to sequences with the release type “Major”) as well as the genomes in the Uncultivated Bacteria and Archaea (UBA) dataset (38). Both datasets contain a high number of incompletely sequenced chromosomes and chromosomal fragments. After discarding sequences present in both databases, we retained 25,827 *oriC* sequences from 1,217 different gammaproteobacterial species, most of which were not identified yet. These sequences constitute the first version of BOriS DB and are available for download at boris.heiderlab.de.

### Comparison to DoriC

Due to the fact that only very few *oriC* sequences are experimentally confirmed for Gammaproteobacteria, and there is no established *oriC* benchmark dataset, a direct comparison of *oriC* identification tools is infeasible. Therefore we compared sequences collected using *γ*BOriS to the current state-of-the-art *oriC* database, DoriC (21, 22). To this end, we created an *oriC* dataset by using the 462 chromosomes present in DoriC as input for *γ*BOriS.

For 330 of the chromosomes listed in DoriC we find the same *oriC* sequence, however, for 156 there is disagreement. To evaluate which of the datasets is more consistent, we calculated pairwise similarity matrices for all sequences in DoriC and the results from *γ*BOriS, respectively. The underlying assumption behind this method is that the *oriC* sequences of different organisms are related evolutionarily and therefore show a high amount of similarity; mis-identified sequences will be more different to the other sequences in the dataset.

Visualization using multidimensional scaling shows that the sequences identified using *γ*BOriS, generally form tighter clusters than the sequences stored in DoriC (fig. 3), which indicates more consistency. This result is also supported by phylogenetic trees derived from the pairwise distance matrices (see supplementary information). A closer inspection of these results shows that while for most orders, *γ*BOriS is more consistent than DoriC (as, e.g., for Vibrionales and Xanthomonadales), the contrary is true for chromosomes from, e.g., Methylococcales and Thiotrichales. For many Gammaproteobacterial endosymbionts, *oriC* sequences are not well-identified neither in DoriC nor by *γ*BOriS, which is due to the fact that most of these genomes lack a *dnaA* gene and rely on a different initiation method (43).

**Fig. 3.**
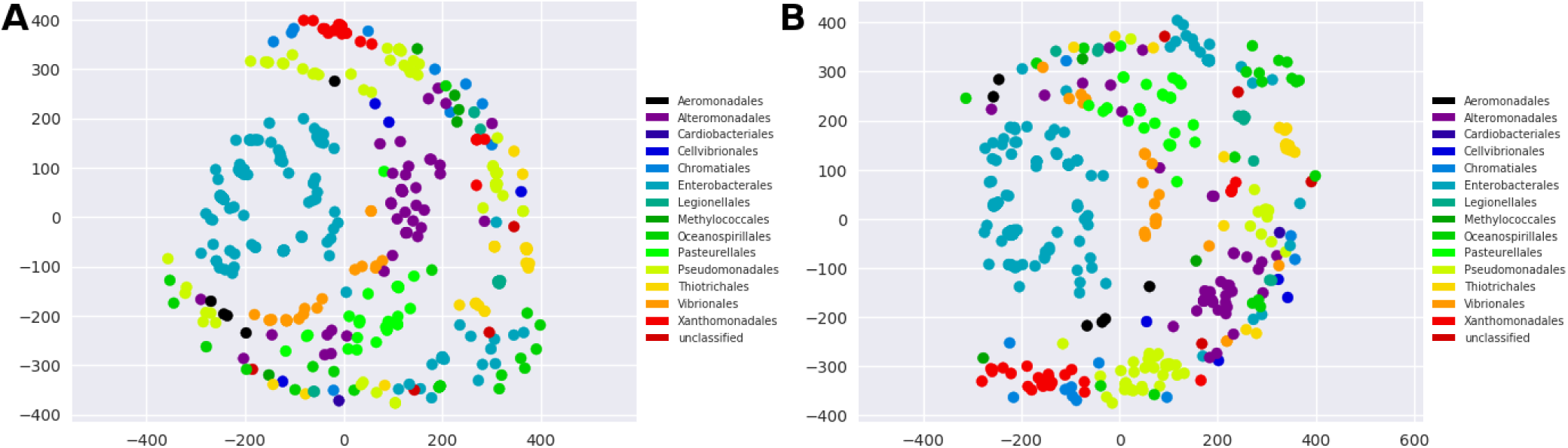
Consistency of the sequences present in *oriC* databases. Multidimensional scaling representation of the distance matrix calculated from all-vs-all pairwise sequence alignments of sequences from *oriC* databases; **(A)** *γ*BOriS (used on chromosomes that DoriC is created from) and **(B)** DoriC.

## Discussion

Currently, one of the most promosing applications for machine learning methods in bioinformatics is the classification and identification of DNA sequences (44, 45). While machine learning methods have already been employed for the identification of origins of replication in yeast (46), *oriC* identification in bacterial chromosomes is still performed based on chromosome-wide nucleotide disparities such as the GC-skew. As these methods are limited to fully sequenced chromosomes, no *oriC* sequences can be identified for a huge number of only fragmentarily sequenced genomes. Furthermore, the methods developed for eukaryotic chromosomes cannot easily applied to bacterial chromosomes as the composition of these sequences are radically different (47). In contrast, *γ*BOriS, which we introduce here and which makes use of a motif-based machine learning method, is able to identify *oriC* sequences on chromosomal fragments as well as full chromosomes of Gammaproteobacteria.

Due to the fact that there is a high degree of variance in *oriC* structure between taxonomic classes (7, 43), we limited the scope of *γ*BOriS to Gammaproteobacteria. Furthermore, as most secondary chromosomes do not rely on the initiator protein DnaA for replication initiation (48, 49), and as they are rather rare in bacterial cells (3), we also excluded these from the scope of the tool and focused only on primary chromosomes.

Using different training datasets, the general approach of *γ*BOriS can easily be adapted for other groups of organisms. Suitable datasets, however, are currently not easily available in the necessary amount and quality (e.g., same-sized, centered, and co-oriented) because current *oriC* identification methods do not provide the identified sequences in this manner. The semi-automatic method used to create an initial *oriC* dataset in this study assumes that (I) *oriC* is intergenic, (II) close to the global GC skew minimum, and (III) defined by the DUE, as well as (IV) the presence of DnaA boxes. The fact that this method requires manual decision-making makes it hard to automate it, but also ensures that the weight of the assumptions can be balanced and adjusted for every single case. Therefore, we consider this method highly accurate, which is supported by the fact that *oriC* sequences identified have been confirmed experimentally (9, 35, 36). Being solely trained on sequences gathered with the method used in this paper, *γ*BOriS can be used as an automatization and extention of it.

By applying *γ*BOriS on fragments deposited in public sequence databases, we created BOriS DB, the largest availabe database of *oriC* sequences to date. BOriS DB currently contains 25,827 sequences from 1,217 species, most of which were not identified yet. The sequences in this database show a high degree of consistency, which indicates a high degree of accuracy in prediction (see fig. 3 and supplementary information). A comparison of BOriS DB to DoriC suggests that both databases are more reliable for some taxonomic groups than for others, with *γ*BOriS, in total, showing a better performance.

*γ*BOriS enables researchers to identify *oriC* sequences on fragments of bacterial chromosomes making it possible to integrate it into next-generation sequencing pipelines. As can be seen from the construction of BOriS DB, this leads to a large amount of newly-identified *oriC* sequences, among which are sequences from organisms that are notoriously hard to sequence and impossible to culture. This enables the use of data-intensive cutting-edge methods such as deep learning (50) for the identification previously unknown initiation factors, which might, due to their high degree of taxonomic specificity, be good candidates for targets of new antibiotics (51). Furthermore, a deeper knowledge of the components of of *oriC* will make it possible to *de novo* design chromosomes with desired replication characteristics and synthetic *oriC* sequences (52). *γ*BOriS as a web tool as well as a stand-alone software, its source code, and BOriS DB are freely available at BOriS.heiderlab.de.

## Conflict of Interest Statement

The authors declare that the research was conducted in the absence of any commercial or financial relationships that could be construed as a potential conflict of interest.

## Author Contributions

TS conceived the project and drafted the manuscript. TS and LM designed *γ*BOriS and performed the machine learning analyses. LM implemented *γ*BOriS. CW provided the ground truth data. RM implemented the web server for *γ*BOriS. TS, TW, CW, and DH discussed the results. DH supervised the project and revised the manuscript.

## Funding

This work was partially funded by the Philipps-University of Marburg. Publication costs for this manuscript were sponsored by the Philipps-University of Marburg.

## Acknowledgements

Calculations on the MaRC2 high performance computer of the University of Marburg were conducted for this research. We would like to thank Mr. Sitt of HPC-Hessen, funded by the State Ministry of Higher Education, Research and the Arts, for installation and maintenance of software on the MaRC2 high performance computer.

## Supplemental Data

Supplemental file 1. Contains more detailed description and discussion of methods and results, Tables S1-S2 and Figures S1–S4.

