## Supplementary file for "γBOriS: Identification of Origins of Replication in Gammaproteobacteria using Motif-based Machine Learning"

### $\gamma$BOriS: Identification of Origins of Replication in Gammaproteobacteria using Motif-based Machine Learning
#### *Supplementary Information*

Theodor Sperlea[1], Lea Muth[1], Christoph Weigel[2], Thorsten Waldminghaus[3] and Dominik Heider[1]

1. Faculty of Mathematics and Computer Science, Philipps-Universität Marburg, D-35043 Marburg, Germany
2. Department of Life Science Engineering, Fachbereich 2, HTW Berlin, D-10318 Berlin, Germany
3. LOEWE Center for Synthetic Microbiology (SYNMIKRO), Philipps-Universität Marburg, D-35043 Marburg, Germany

February 22, 2019

**Abstract**

The biology of bacterial cells is, in general, grounded on the information encoded on circular chromosomes. Consequently, maintenance of these through regulation of chromosome replication is an essential process which mostly takes place at a single genomic site, called the origin of replication (*oriC*). To enable the systematic study of *oriC* function with cutting-edge methods and the potential to identify novel antibiotic drug targets, high numbers of correctly identified sequences are needed. However, current methods for identyfing *oriC* sequences rely on chromosome-wide nucleotide disparities and are therefore limited to fully sequenced genomes, leaving a superabundance of genomic fragments in public databases unstudied.

Here, we present $\gamma$BOriS (Gammaproteobacterial *oriC* Searcher), which is able to identify *oriC* sequences on gammaproteobacterial chromosomal fragments as well as on full chromosomes at a high accuracy by employing motif-based DNA classification methods. Using $\gamma$BOriS, we created BOriS DB, which currently contains 25827 *oriC* sequences from 1217 species, thus making it the largest available database for *oriC* sequences to date.
$\gamma$BOriS as a web tool as well as a stand-alone software, its source code, and BOriS DB are freely available at `BOriS.heiderlab.de`. **Contact:**

### 1 Results and Discussion

#### 1.1 Dataset generation

The sequence datasets used in this study are based on an initial dataset of 565 *oriC* sequences identified in Gammaroteobacterial chromosomes using the method first described in [4] and subsequently used in [9, 10]. This dataset was chosen because the sequences in it were of equal size, co-centered and co-oriented, which is not the case for automatically generated *oriC* datasets but necessary for most machine learning methods. Furthermore, this dataset contains only Gammaproteobacterial *oriC* sequences to reduce variation. Note that, for some organisms, as e.G. from the genus Pseudomonas, this dataset contains two *oriC* sequences, which might act as bipartite or independent *oriC* [7, 11]. See figure S1 for detailed taxonomic distribution. In order to generate positive datasets, these sequences were enlargened and shortened to sequences ranging from 500 bp to 1500 bp in steps of 50 bp.

From this initial dataset, we derived a set of unique seed sequences by extracting the central 9 bp motif from every single sequence, which, in many *oriC* sequences represents a DnaA box, i.e. a binding site for DnaA.

A negative dataset (i.e. a dataset consisting of non-*oriC* sequences) was assembled by choosing one sequence from every chromosome represented in the positive dataset that is centered around the same or a similar seed sequence (up to 2 mismatches, increasing, unless a better match is present) as the corresponding sequence in the positive dataset. As for the positive dataset, negative dataset sequence lengths were varied in a range of between 100 and 1500 bp, in steps of 50 bp.

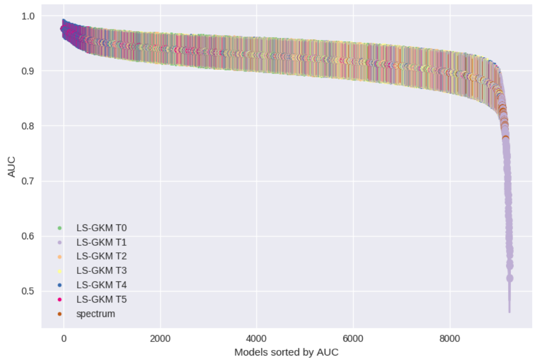


(a)

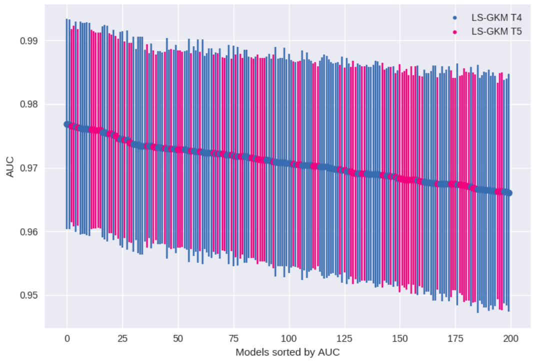


(b)

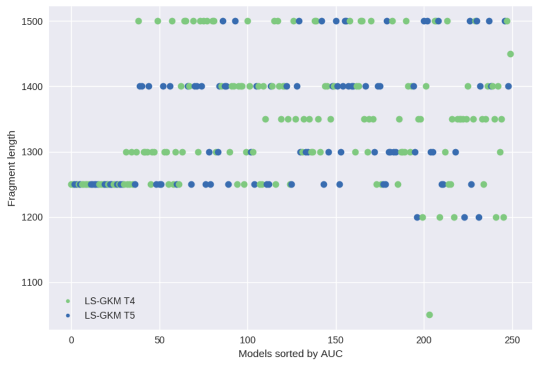


(c)

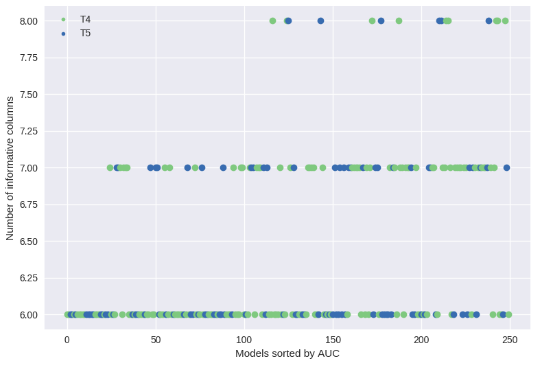


(d)

Figure S1: **Comparison of the *oriC* classification performance of different k-mer SVMs**. **(a)** Visualization of the performance of all models. Error bars represent confidence intervals calculated using a permutation test. **(b)** Visualization of the performance of the 200 best models as measured by AUC. Error bars represent confidence intervals calculated using a permutation test. **(c)** Visualization of the fragment length the 250 best models (as measured by AUC) were trained on. **(c)** Visualization of values of the number of informative columns that was chosen as parameter for the 250 best models (as measured by AUC).

#### 1.2 *k*-mer SVM training and evaluation

Models from the LS-GKM package [12] as well as Spectrum Kernel models [5] were trained using 70% of the negative and positive *oriC* sequence datasets while examining all possible combinations of sequence sizes and parameter variations, totalling around 13000 models (see tab. S1 for details). The models were evaluated using the

Table S1: Parameters of the machine learning models used here and the ranges of their values. Ranges are presented in square brackets that contain the start and end value and step size. Values in parentheses suggest a list of possible values.

| Parameter Name | Number LS-GKM | Spectrum |
|---|---|---|
| Fragment Length | [100, 1500, 50] | [100, 1500, 50] |
| Motif length | [6, 12, 1] | [6, 12, 1] |
| Model type | (T0, T1, T2, T3, T4, T5) | - |
| Informative Columns | [5, 11, 1] | - |
| Allowed Mismatches | [1, 4, 1] | - |
| Total amount of models | 12877 | 209 |

remaining 30% of the datasets. Almost all models achieved an AUC of larger than 0.8 (see fig. S2a). Due to this and the number of models trained, the significance of performance difference as calculated using the DeLong test [3] is not informative.

A more detailed look at the parameters used for the best performing models shows that these are dominated by LS-GKM models with the type T4 and T5 (see fig. S2b). Furthermore, almost all of the 25 best performing models were trained using a fragment length of 1250 bp and 6 informative columns (see tab. S2), while there is more variation in these parameters for the 250 best performing models (see figs. S2c and S2d). The other parameters provided by LS-GKM, i.e. the word length and number of mismatches, seem not to have a large impact on model performance as they show a high amount of variation in the best performing models (see tab. S2).

#### 1.3 Implementation of *oriC* identification method

##### 1.3.1 Turning a classifier into an identifier

In order to transform a DNA classification model into a model that is able to identify a genomic position where the classification value is maximal, we see three different possible approaches:

1) LS-GKM models are able to use DNA fragments as inputs that have different lengths from the DNA fragments they were trained on. Therefore, one could split a large-scale DNA sequence into a few very large DNA fragments, score these using a LS-GKM model, then splitting the highest scoring one into subfragments and scoring these, and so on, until having reached the target fragment size. However, we were not able to prove that the fragment containing *oriC* will always have the

Table S2: Parameters and evaluation results of the 25 SVM models with the highest AUC values.

| Index | Type | Fragment length | word length | informative columns | mismatches | AUC | DeLong error |
|---|---|---|---|---|---|---|---|
| 1 | LS-GKM T4 | 1250 | 10 | 6 | 4 | 0.97685 | 0.01649 |
| 2 | LS-GKM T4 | 1250 | 10 | 6 | 3 | 0.97678 | 0.01647 |
| 3 | LS-GKM T5 | 1250 | 10 | 6 | 3 | 0.97657 | 0.01515 |
| 4 | LS-GKM T5 | 1250 | 8 | 6 | 2 | 0.97654 | 0.01576 |
| 5 | LS-GKM T4 | 1250 | 12 | 6 | 4 | 0.97647 | 0.01653 |
| 6 | LS-GKM T5 | 1250 | 10 | 6 | 4 | 0.9763 | 0.01543 |
| 7 | LS-GKM T4 | 1250 | 11 | 6 | 4 | 0.97626 | 0.01672 |
| 8 | LS-GKM T4 | 1250 | 9 | 6 | 3 | 0.97616 | 0.01655 |
| 9 | LS-GKM T4 | 1250 | 9 | 6 | 2 | 0.97616 | 0.01655 |
| 10 | LS-GKM T4 | 1250 | 8 | 6 | 2 | 0.97609 | 0.01675 |
| 11 | LS-GKM T4 | 1250 | 11 | 6 | 3 | 0.97602 | 0.01674 |
| 12 | LS-GKM T5 | 1250 | 7 | 6 | 1 | 0.97599 | 0.01563 |
| 13 | LS-GKM T5 | 1250 | 8 | 6 | 1 | 0.97595 | 0.01551 |
| 14 | LS-GKM T5 | 1250 | 9 | 6 | 3 | 0.97588 | 0.01539 |
| 15 | LS-GKM T5 | 1250 | 10 | 6 | 2 | 0.97588 | 0.0153 |
| 16 | LS-GKM T5 | 1250 | 9 | 6 | 2 | 0.97588 | 0.01539 |
| 17 | LS-GKM T4 | 1250 | 8 | 6 | 1 | 0.97561 | 0.01658 |
| 18 | LS-GKM T4 | 1250 | 10 | 6 | 2 | 0.97557 | 0.01685 |
| 19 | LS-GKM T4 | 1250 | 12 | 6 | 3 | 0.9754 | 0.01696 |
| 20 | LS-GKM T5 | 1250 | 12 | 6 | 4 | 0.97536 | 0.01571 |
| 21 | LS-GKM T5 | 1250 | 11 | 6 | 4 | 0.97533 | 0.01562 |
| 22 | LS-GKM T4 | 1250 | 7 | 6 | 1 | 0.97512 | 0.01649 |
| 23 | LS-GKM T5 | 1250 | 11 | 6 | 3 | 0.97505 | 0.01568 |
| 24 | LS-GKM T5 | 1250 | 12 | 6 | 3 | 0.97457 | 0.01569 |
| 25 | LS-GKM T4 | 1250 | 9 | 7 | 2 | 0.97453 | 0.01681 |

highest classification score, independent of fragment size (i.e. size of "noise" DNA sequences). Therefore, this approach is not reliable.

2) In order to create fragments of the size that the LS-GKM model was trained on, one could use a moving window approach, starting each candidate fragment a certain step size after the start point of the last fragment. For this approach to work, one would need to identify the highest step size for which the classifier still identifies each *oriC* sequence. However, this approach would lead to a number of LS-GKM scoring steps that grows linear with input sequence size, which, in the case of full bacterial chromosomes, will most likely be in the range of millions. Therefore, this approach is not practical for our application.

3) In order to choose a reduced number of candidate fragments, one might employ a "external" method, that is not based on the LS-GKM model itself. Reducing the number of candidates leads, of course, to a lower coverage of the input sequence, which, in turn, might lead to high numbers of false negatives, as some positive sequences might not even be identified as candidates.

We decided to employ the third approach, using the seed sequences that were extracted from the initial dataset (see sect. 1.2) in order to create candidate fragments of 1250 bp from input sequences that are centered around any one of the instances of the seed sequences. As the seed sequences that can be extracted from the test sequences (as used in sect. 1.2) are all present in the seed sequences extracted from the training sequences, we conclude that our choice of seed sequences will not lead to many false negatives, which is also supported by the results shown in sect. 1.3.2. Note that the size and composition of these seed sequence list is one of the largest contributors to the runtime of $\gamma$BOriS.

##### 1.3.2 Identification of optimal cutoffs

Using the classifier trained as described in sect. 1.2 to identify a DNA fragment from a larger set of candidate sequences (created as described in sect. 1.3.1) turns the balanced classification problem into a highly inbalanced one. Therefore, cutoffs between positive and negative labels have to be identified that maximize metrics suitable for imbalanced classification problems, such as the F1 score. Furthermore, we decided to make use of the concept of classification with abstaining (creating a range of values in which the classifier abstains from assigning a classification label to candidate sequences) [1]. This leads to an additional aim of the cutoffs, namely minimizing the number of samples for which the classifier abstained. In order to identify a setting that best fulfills both aims, four different classification score normalization methods were investigated combined with different cutoff configurations:

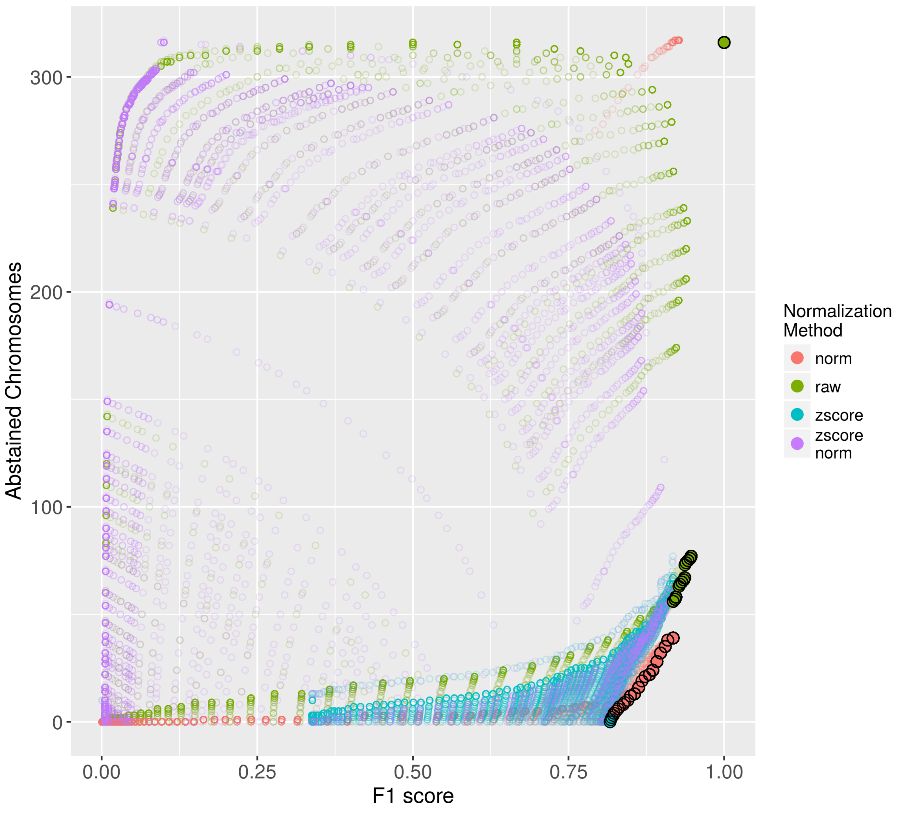


Figure S2: **Relationship between F1 score and abstaining space given different normalization methods on the training dataset.** The number of rejected chromosomes is the number of chromosomes, for which the positive fragment is scored between the two cutoffs and is thus abstained. Every dot represents one set of cutoff parameters and a normalization method; dots with black border represent parameter sets on the pareto front.

- No preprocessing ("raw" in fig. S3).
- Normalizing the classification scores of the fragments gathered from one input sequence to a range of [0, 1] ("norm" in fig. S3).
- Normalizing the classification scores of the fragments gathered from one input sequence by calculating z-scores ("zscore" in fig. S3) using the formula

$$z = \frac{(X - \mu)}{\sigma}$$

where $x$ is the classification score of a fragment, $\mu$ is the mean of the classification scores of the fragments of one input sequence and $\sigma$ is the standard deviation of the classification scores of the fragments of one input sequence.

- Normalizing the z-score normalized classification scores of the fragments gathered from one input sequence to a range of [0, 1] (”zscore norm” in fig. S3)

Using the same train-test split as in sect. 1.2, we identified all cutoff-normalization configurations that are pareto optimal in maximizing the F1 score while minimizing the number of rejected chromosomes using the R package rPref. From these, we decided to use the cutoff-normalization configuration for $\gamma$BOriS that creates no false negative results on test data (as well as the training data), while taking a higher number of false positive decisions into account. This way, researchers using $\gamma$BOriS will be presented some wrong fragments, but no fragments that do contain *oriC* will be among the discarded sequences. Therefore, we chose a lower cutoff of 0.41 and a upper cutoff of 0.99 for data that were normalized to a range between 0 and 1, which resulted in a F1 score of 0.94 and 11 rejected chromosomes on test data.

##### 1.3.3 Comparison to DoriC

The current state-of-the-art method for the identification of *oriC* sequences is Ori-Finder, which makes use of the Z-curve method and is, thus, only useable for fully sequenced chromosomes [8, 13]. Due to the fact that, currently, there is no *oriC* benchmark dataset, it is impossible to compare our method to Ori-Finder. Therefore, we decided to compare DoriC (version 5) [6], which was created using Ori-Finder, to the results of using $\gamma$BOriS on the same chromosomes Ori-Finder was used on to create DoriC. The comparison between the datasets is based on the similarity of the sequences identified as *oriC* sequences in the respective datasets, thus, their internal consistency. The reasoning behind this method of comparison is that *oriC* sequences from closely related organisms are most probably rather similar; therefore, a database that contains *oriC* sequences that are more consistent in taxonomic subgroups must have been created using a more precise method for *oriC* identification.

Consistency of databases was evaluated using pairwise global alignments (performed using the align submodule from Biopython [2]) for all sequences in both DoriC and the respective results created using $\gamma$BOriS. An overview over these values is given in fig. 2 of the main paper after applying multidimensional scaling and in fig. S4, for which a dendrogram was created from the similarity matrices. Both visualizations show that the sequences resulting from $\gamma$BOriS are more consistent,

Table S3: Mean and standard deviation of the scores of pairwise global alignments in taxonomic groups as an indication of consistency. A higher score indicates a higher degree of similary.

| Taxon | BOriS DB | | DoriC | |
|---|---|---|---|---|
| | Mean | St.dev. | Mean | St.dev. |
| Aeromonadales | 1032.16 | 146.23 | 977.33 | 158.6 |
| Alteromonadales | 912.01 | 352.23 | 1119.65 | 130.35 |
| Cellvibrionales | 927.5 | 194.55 | 910.75 | 202.47 |
| Chromatiales | 861.74 | 137.19 | 851.64 | 134.12 |
| Enterobacterales | 982.66 | 149.34 | 977.19 | 150.67 |
| Legionellales | 1005.09 | 211.07 | 983.37 | 203.7 |
| Methylococcales | 940.64 | 189.9 | 969.11 | 218.48 |
| Oceanospirillales | 823.4 | 145.01 | 821.46 | 137.68 |
| Pasteurellales | 947.3 | 105.91 | 920.69 | 106.99 |
| Pseudomonadales | 849.24 | 117.9 | 886.47 | 132.95 |
| Thiotrichales | 909.72 | 156.8 | 985.57 | 185.74 |
| Vibrionales | 1006.08 | 118.29 | 927.43 | 127.3 |
| Xanthomonadales | 987.11 | 140.4 | 910.18 | 140.55 |

as sequences from the same taxonomic subgroup show a higher degree of clustering than those from DoriC. To quantify this, we calculated average sequence similarities for taxonomic orders (see tab. S3). Except for Alteromonadales, Methylococcales, Pseudomonadales and Thiotrichales, the sequences gained using $\gamma$BOriS show higher mean similarities.

The performance of $\gamma$BOriS is, most likely, due to low numbers of organisms from these groups in the initial dataset (especially for Thiotrichales and Methylococcales). Chromosomes of Pseudomonadales are known to contain two *oriC*, whose function is not fully clear yet (Are both needed for correct initiation or can initiation occur at both? Do the loci need to interact for initiation to occur?) [11, 7]. In the ground truth dataset used for training and testing of $\gamma$BOriS, both sequences are present; however, only one of the sequences is usually identified as *oriC* by $\gamma$BOriS (see fig. S1). This might reflect a difference in function of these two *oriC* sequences, but further research into both Pseudomonas *oriC* as well as the decision-making $\gamma$BOriS would be needed to make this claim.

[7] Feng Gao. Bacteria may have multiple replication origins. *Front. Microbiol.*, 6, apr 2015.

[13] Hao Luo, Chun-Lan Quan, Chong Peng, and Feng Gao. Recent development of ori-finder system and DoriC database for microbial replication origins. *Briefings in Bioinformatics*, jan 2018.

[14] Donovan H. Parks, Christian Rinke, Maria Chuvochina, Pierre-Alain Chaumeil, Ben J. Woodcroft, Paul N. Evans, Philip Hugenholtz, and Gene W. Tyson. Recovery of nearly 8,000 metagenome-assembled genomes substantially expands the tree of life. *Nature Microbiology*, sep 2017.

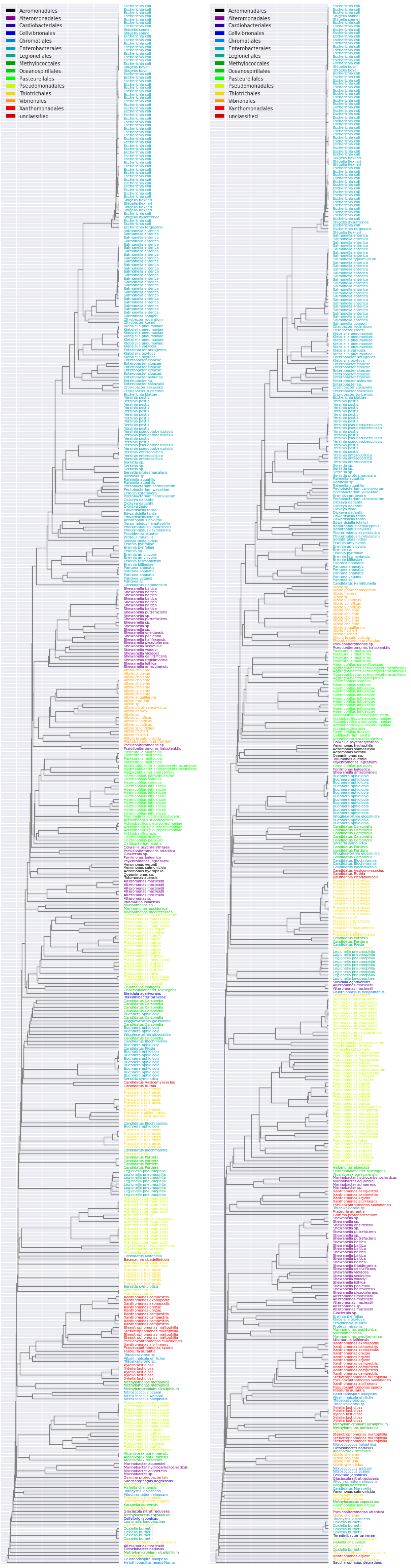


(a) (b)

Figure S3: **Similarity of sequences present in *oriC* databases**. Dendrograms were created from similarity values calculated from global pairwise alignments. a) Sequences identified using $\gamma$BOriS in the chromosomes that were also used as input for Ori-Finder for the creation of DoriC, b) DoriC (version 5.0).
